# Multiple Xpa In vivo crystallization routes in HEK 293 human cells

**DOI:** 10.1101/2025.02.12.637855

**Authors:** O. Leymarie, A. Skobelkina, P. Montaville, C. Brewee, T. Isabet, H. Chauvet, E. Pereiro, L. Chavas, F. Jamme

## Abstract

Phase changes of macromolecules in living cells gained recently a major interest in cell biology as liquid liquid phase separation or liquid solid phase transition phenomenon being observed in increasing number of biological or pathological processes. Comprehensive characterization of these phases remains challenging and requires new methodological approaches for their complete description across spatial and temporal scales. We propose a combination of imaging methods applied to a model system and report the *in vivo* crystallization pathways of a macromolecule from its solution to crystalline states at the cell population level down to the meso scale. Combining various live fluorescence based techniques and a high resolution cryo imaging technique within unaltered cryopreserved cells, we could described the unexpectedly wide landscape of the *in vivo* crystalline states of the fluorescent coral derived protein xpa, gain information of crystal growth dynamics *in cellulo*, and provide hypothesis of the crystal nucleation requirements of this stochastic process.

## Introduction

*In vivo* crystallization of biological molecules is a specific and rare phenomenon that has been described throughout the living kingdom ^1,2^. It has been reported for physiologically relevant processes (storage in plants, catalysts, such as crystalloid cores within peroxysomes, in yeast and eucaryotes, protists, protection and stabilization in viruses and prokaryotes for example), related to pathological conditions (examples: immunoglobulinemia, Nemaline myopathy, Charcot Leyden disease, cataract, hemoglobin C disease, Wilson’s disease and some tumors (Reinke crystals) and also for recombinant artificial systems as an alternative approach to classical *in vitro* crystallogenesis ^3–6^ . Crystals produced in vivo have been used for 3D structure determination, and the method holds a number of advantages as compared to the well established semi-empirical in vitro crystallization methods currently in use. However, *In vivo* crystallization of a protein for macromolecular structure determination in the highly heterogeneous environment of a living cell could be counterintuitive at first sight. This is why understanding *in vivo* crystallization key parameters is essential for this approach to be an advantageous alternative to *in vitro* crystallization of purified macromolecules for which semi empirical screening remains the major bottleneck. So far only a handful set of *in vivo* grown protein crystals have led to high resolution structures ^2^, and efforts have been made to characterize and understand the mechanisms underlying this process to establish the potential of *in vivo* crystallogenesis for the structural biology field. The rationalization of in vivo crystallization understanding and mastering lies on the requirement of a multiscale approach from the cell population to characterize variability of the process down to the meso scale at which nucleation processes occur. A multiscale approach revealed to be also crucial since the final size of *in vivo* grown crystals spans length scales from the tens of nanometers to the tens/hundreds of micrometers ^2^ . If electron microscopy has been used ^2^ to describe *in vivo* grown crystals in their native environment, gathering statistics from this technique remains difficult and should be used when the system is comprehensively described at wider scales. This is particularly true for crystal nucleation sites within the cellular environment that are of a tremendous interest to the field and remains poorly understood. For this purpose, a dynamical view of crystal growth is required to assign within cellular substructures the chances of crystal nucleation as well as the expected morphology of the fully-grown crystal that might impact its diffraction property. We have consequently designed a multiscale workflow comprising multiphoton microscopy, confocal microscopy, DIC microscopy and correlated cryo fluorescence and soft X-ray microscopy have been chosen to cross the spatial scales and multiphoton microscopy associated to spinning disk confocal microscopy were used for dynamics evaluation of crystal growth. As a study model, we chose the coral-derived fluorescent protein variant Xpa that has been shown to crystallize in human embryonic kidney 293 (HEK293) cells during the semi-rational engineering of a protein with higher optical properties ^7–9^. Interestingly, Tsutsui *et al*. showed that an *in cellulo* diffraction data collection, on a single crystal (10µm of length), allowed for structure determination of *in vivo*-grown Xpa crystal. Xpa exhibits the ability to crystallize in the nucleus as well as in the highly heterogeneous cytoplasm, addressing the role of the impact of the environment complexity to Xpa crystal formation and growth. Xpa has been shown to crystallize in various types of cells ^6,9^ but also, more recently in *C. elegans* ^10^. In all cases found in the literature, *in vivo* Xpa crystallization has been shown to be a stochastic event within a small subset of cells (roughly 0.1% of cells^9^), which make investigation at the cell population level essential. In the present work, we chose to use the specific HEK293 Free Style cell line, due to its heterologous expression properties, to validate our multiscale and multimodal imaging approach and test its robustness for other *in vivo* grown crystallization systems. This work allowed us to record a comprehensive atlas of *in vivo* grown Xpa crystals at unprecedented levels of detail, providing some information on Xpa crystal growth dynamics, and hints for Xpa crystal nucleation sites requirements within living cells.

## Results

### 1 Xpa crystallization in HEK293 Free Style cell type as a suitable model for protein *in vivo* grown study

To get insight into the determinants of Xpa *in vivo* crystallization at the meso scale, the HEK293 Free Style (HEK293FS) cell line was chosen for its handling convenience, its high heterologous protein expression ability, and its spherical shape (that might impact the Xpa crystal growth and ultimate shape) while maintained in suspension. To confirm the suitability of this expression system for *in vivo* crystallization we used multiphoton microscopy (fluorescence^11,12^ and SHG^13–15^) to monitor heterologous expression levels via Xpa fluorescence as well as its *in vivo* crystallizability at the cell population level (Fig. 1A and Supplementary Fig. 1). The expression level of Xpa in HEK293FS cells shows a significant heterogeneity from cell to cell, as evidenced by the variable fluorescence intensity over the whole population (Fig. 1A). The time course of Xpa expression was followed, as early as 3 hours after transfection (Supplementary Fig. 1). Cells exhibit a linear increase of the fluorescence level (see Supplementary Fig. 1A right panel) within the first hours before reaching a plateau 1 day post transfection. From 1 to 5 days post transfection no overall fluorescence level increase was observed throughout the cell population (data not shown).

**Figure 1.**
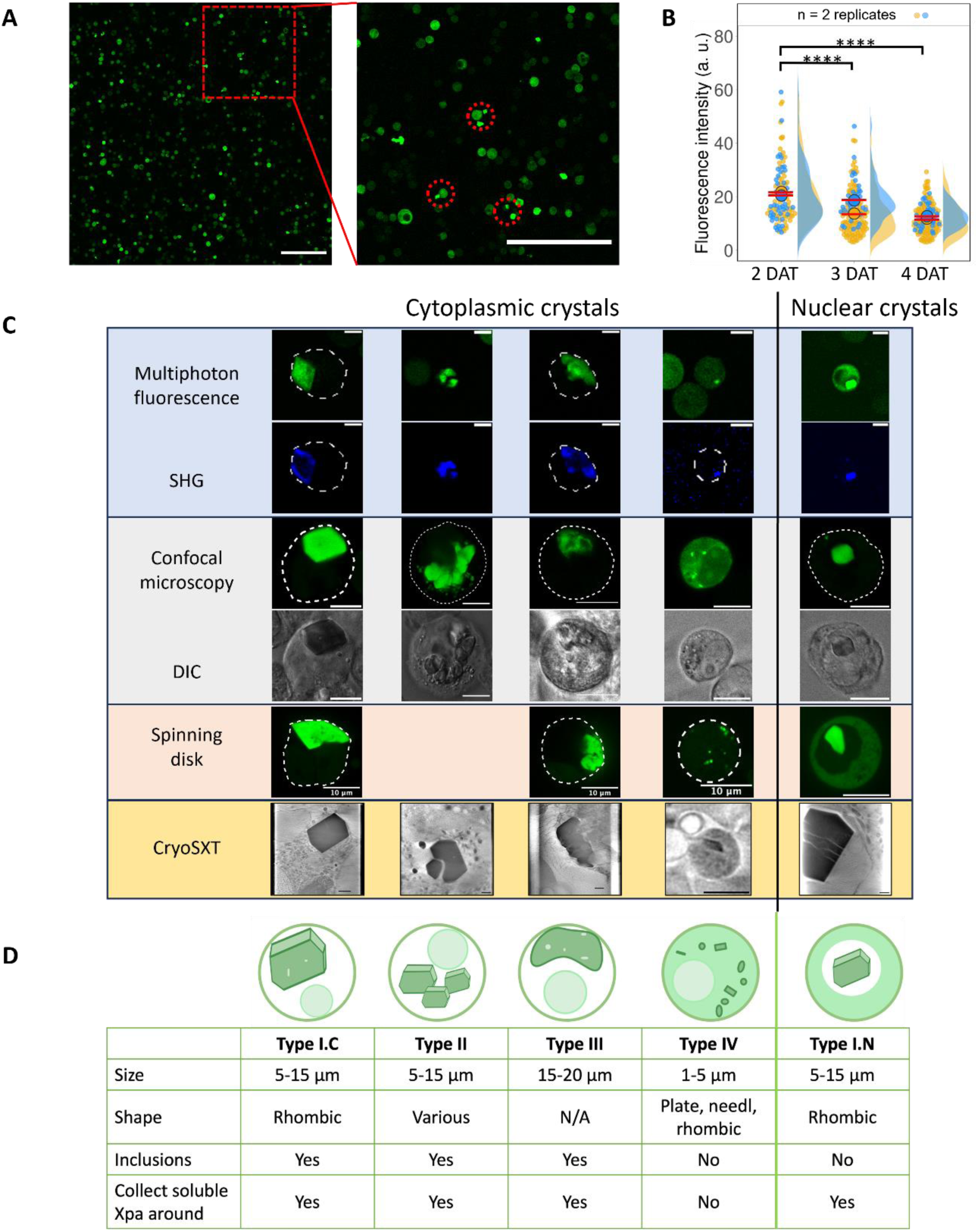
Crystals atlas. **(A)** Representative multiphoton fluorescence images obtained by multiphoton microscopy after 3 days after transfection (DAT) of XPA. Scale bars: 200 µm. **(B)** Change in fluorescence intensity as a function of time after transfection. P-values are lower than 0.0001. **(C)** Classification of different types of crystals obtained by multiphoton fluorescence microscopy, SHG microscopy, confocal microscopy, DIC microscopy, spinning disk microscopy and cryoSXT. **(D)** Atlas of crystal types with key parameters.

Three days post transfection, *in vivo* crystals of Xpa could be readily observed in HEK293FS cells maintained in suspension (Fig. 1A). Crystallization events were detected mainly in the cytoplasm, but also within the nucleus despite a lower occurrence as previously described^9^. In almost all cases (see below), Xpa crystallization is followed by the depletion of all soluble Xpa in the respective compartments (Fig. 1C columns 1, 2, 3, 5 and Supplementary Fig. 1E). As previously described^9^, after Xpa crystallization and the recruitment of all soluble Xpa within the solid crystalline phase, no soluble Xpa diffusion was observed, suggesting that the dilute protein does not cross the nuclear membrane either ways.

Post-plateau, among cells that did not experienced Xpa crystallization, a steady decrease of the number of highly fluorescent cells (between two and four days post transcription) was observed (Fig. 1B). Furthermore, crystallization of Xpa seemed favored in cells smaller than 15µm in diameter (below 250 µm^2^) and crystallization in larger cells appeared to occur at a later stage of the experiment (Supplementary Fig. 1B). The evolution of the fluorescence intensity distribution correlated to the cell size over time showed that the 15 µm diameter cells exhibited the highest Xpa densities and were more favorable to Xpa crystallization than smaller or larger cells (Supplementary Fig. 1B -). This observation suggests that the *in vivo* crystallization process of Xpa is facilitated by high intracellular concentrations of the protein. Note also that the smaller number of intranuclear crystals compared to the intracytoplasmic crystals could be partly due to the lower nuclear Xpa concentration than its cytoplasmic counterpart.

Similarly to the case of Xpa crystallization in HEK293 adherent cells^9^, the first *in vivo* grown crystals were observed within the first day after transfection and the amount of crystals appearing within the cell population increased from 1% of Xpa expressing cells, at two days after transfection, up to 18% , 4 days after transfection (Supplementary Fig. 1C and Supplementary Fig. 2). Conversely, Xpa crystallization is 200 times more frequent in the HEK293FS cells than what has been reported for the HEK293 adherent cells. In summary, the HEK2943FS model reproduces what has been previously observed, but with an increased Xpa nucleation occurrence and revealed to be a suitable model to explore and characterize exhaustively *in vivo* protein nucleation and crystal growth modalities at various spatial and time scales.

### 2 Multiscale morphology variability description of *in vivo* grown Xpa crystals –

Taking advantage of a combined multitechnical approach, an atlas of the *in vivo* grown Xpa crystals at the cellular and subcellular levels can be established (Fig. 1C). Multiphoton fluorescence and SHG microscopy provide global and local information on the *in vivo* grown Xpa dense solid phases, such as their mono or polycrystalline nature as well as the spatial organization of their crystal domains. However, the low sensitivity of the multiphoton based fluorescence and its limited confocality, considering the size of the objects of interest, impaired the possibility of identifying small local fluorescence intensity variations that can be related to local Xpa concentration in solution or inside minute crystals. To tackle this issue, we can take advantage of the one-photon fluorescence microscopy and the cryo-soft X-ray tomography (cryoSXT)^16–21^, supplemented, when required, by cryo-structured illumination microscopy (cryoSIM)^22^ correlative imaging. These techniques can offer a finer description of the Xpa density within the crystals, give information on local soluble Xpa concentration variations, and provide a high resolution description of the various *in vivo*-grown Xpa condensates within their native environment. When combined, these techniques provide a descriptive multiscale overview of the various Xpa liquid to solid state transition pathways *in situ*.

At the single cell level, *in vivo* grown Xpa crystals can be classified in five main groups based on their morphology and intrinsic characteristics. The following classification is proposed:

Type I crystals that can be subdivided in two classes: type I.C (C for cytoplasmic) and type I.N (N for nuclear) crystals relate to the previously described Xpa crystals. Type I Xpa crystals are, at the first order, characterized by their regular rhombic shape and uniqueness within the cell (Fig. 1B columns 1 and 5). Their sizes range from 5 to 15µm in their largest length, with a higher size discrepancy in the cytoplasm as opposed to the nucleus environment (see Fig3 series a *vs* series b). While multiple types of crystals are found in cytoplasm of HEK293FS cells, type I crystals is the only type found in in their nucleus. However, if type I.N crystals are highly homogeneous in term of Xpa density (Fig. 1B column 5), those found in the cytoplasm exhibit defects within the crystal core, as seen by the fluorescence inhomogeneity throughout the crystal section (Fig. 1 column 1 spinning disk technique and Fig. 2A first row or top), or on their surface with localized face defects (Fig. 2A second row or middle, Fig. 3D and movie 1). This observation reflects two entirely different environments: the crowded cytoplasm compared to the more homogeneous nucleoplasm. Indeed, cryoSXT tomogram segmentation of type I.C Xpa crystals shows variations of the carbon density (i.e. local decrease of density) within the crystal core whose shape and dimensions are identical to the surrounding lipid droplets and mitochondria (Fig. 2B first row). Noteworthy, type I.C crystals without defects were also observed but remain rare as observed in cryoSXT tomograms (an example in Fig. 1C column 1 cryoSXT cross section). This suggests that Xpa crystals engulfed some intra-cytoplasmic organelles during their growth. Such inner defects were not observed for any nuclear grown crystals, but cracks probably arising from the vitrification process are detected (Fig. 1B row 5 cryoSXT cross section). As described above, these two subtypes of Xpa crystals emerged directly from the structurally different nature of both cellular compartments.

**Figure 2.**
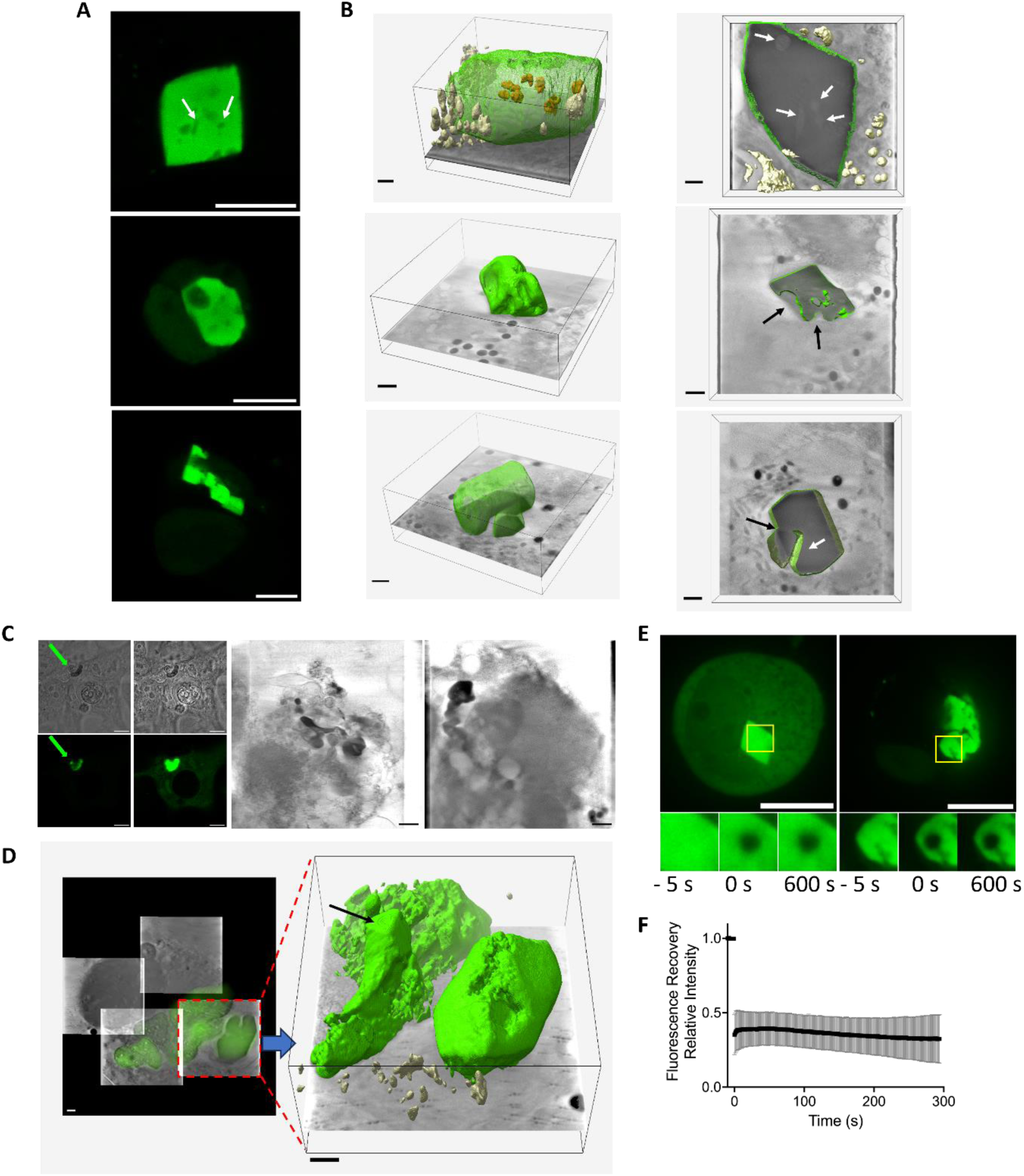
(A) Confocal fluorescence image of a type I.C with inner defects (top), type I.C with surface defects (middle) and type II *in vivo* grown Xpa crystals (scale bar is 10 µm), (B) left column: Segmented volume with a xy 2D slice of 3D reconstructed soft X-ray cryotomograms of similar crystal types as panel A (scale bar is 1 µm) and (B) right column: Segmented volume in the xy plane with a 2D slice of 3D reconstructed soft X-ray cryotomograms shown in the left column (scale bar is 1 µm). Contours of the crystals are shown in green meshes or surfaces, outer vesicules and membranes are shown in beige and engulphed vesicules or mitochondria in the Xpa crystal in brown. White arrows show the internal defects of the Xpa crystals and the black arrows show the surface defects of the Xpa type I.C crystals. For the type III Xpa crystal, the white arrow shows the Xpa depleted region between the two crystals faces, the black arrow shows the fused region between the two crystals. (C) Type III Xpa crystals. Left panel: DIC and confocal fluorescence image desaturated (left) or saturated (right) showing fluorescence distribution inside the Xpa solid condensate (green arrow) within the cell (scale bar 10µm). Right panel: 2D slices from reconstructed cryoSXT tomograms of type III Xpa crystals showing the heterogeneity within type III Xpa crystals (scale bar 1 µm). (D) left panel: mosaic of 4 cryoSXT tomograms 2D planes registered on widefield reconstructed cryofluorescence plane from cryoSIM recorded data of a type III Xpa crystal within a cell (scale bar 1µm). Right: 3D segmentation of on cryoSXT reconstructed tomogram showing a type I.C crystal next to a type III crystal. Both Xpa crystals are coloured in green, carbon dense vesicles and membrane fragments are coloured in beige. The black arrow shows a crystal I.C within the type III crystal (scale bar 1µm). (E) Fluorescence images of representative cells containing type I.C and III crystals analyzed by FRAP. Scale bar: 10 μm. (F) Quantification of the FRAP data from 7 different experiments.

**Figure 3.**
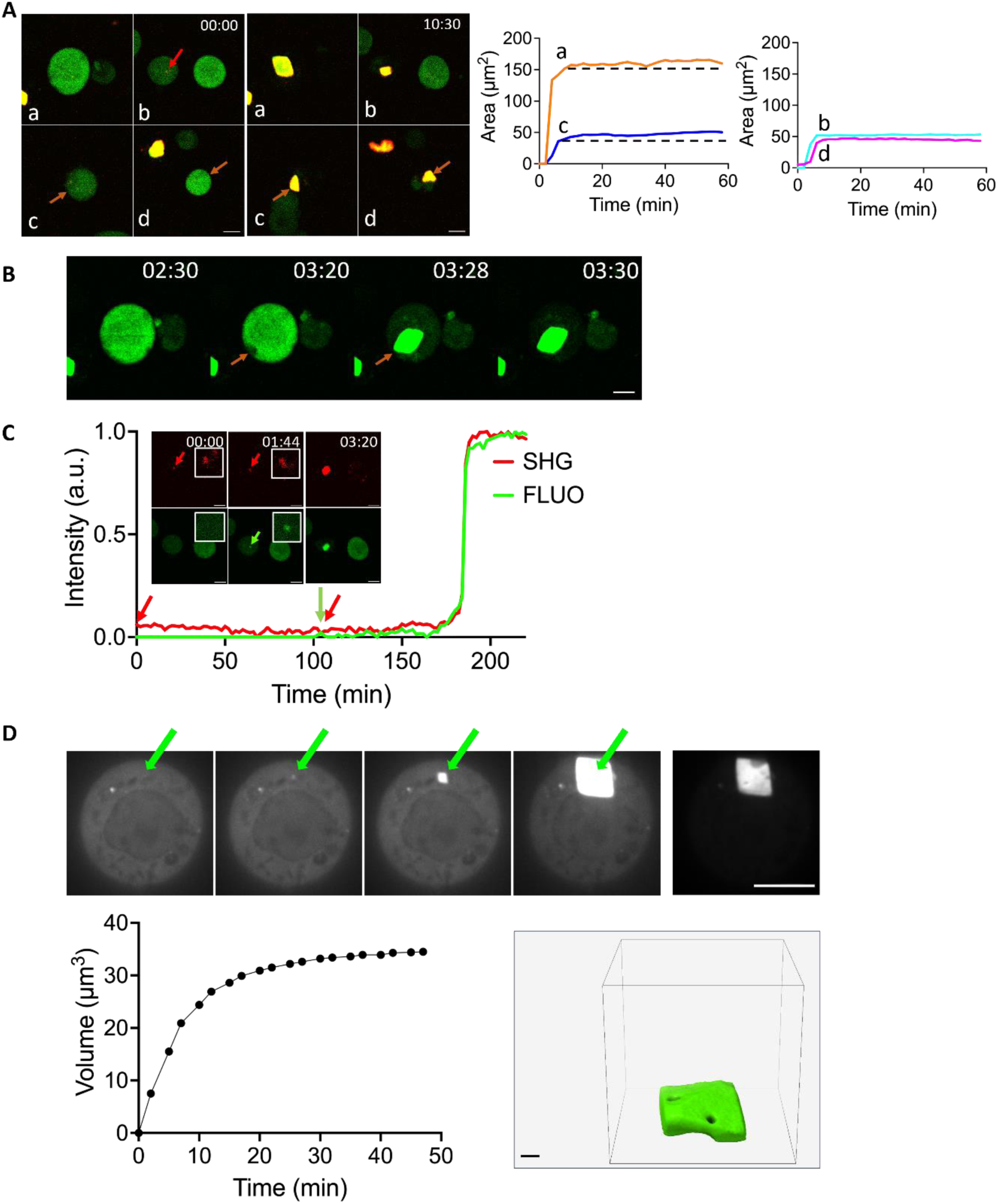
Dynamics of type I.C and type III Xpa crystal growth. (A) left: multiphoton merged fluorescence and SHG channels (green: fluorescence, red: SHG) images of 4 timeseries (type I.C crystal in series a and b and type III crystals in series c and d) at time 00:00 and time 10:30 (hh:mm). Red arrows depict the crystal nucleation areas at time 00:00 for series b to d and for series c and d at time 10:30 corresponding to the plasma membrane of the respective cell (scale bar 10µm). Right : SHG channel derived area of the crystal growth from series a to b over time. Left chart: crystals exhibiting a two phase growth rate, right chart: crystal exhibiting a single phase growth rate. The origin of the time axis has been arbitrarily chosen as the frame before the SHG signal of the crystal appeared. (B) multiphoton fluorescence snapshots of series a type I.C Xpa crystal growth exhibiting plasma membrane alteration preceeding crystal growth (see movie 1 series a). First frame at 02:30 (hh:mm), second frame at 03:20 (8 minutes before crystal appearance), third frame at 03:28 first frame where the crystal is seen, fourth frame 03:30 (crystal in its slow growth rate phase). Red arrows shows the membrane deformation location at 03:20 and 03:28. scale bar 10µm). (C) Relative multiphoton fluorescence (green) and SHG (red) intensity over time of series B type I.C crystal growth. The red arrow shows the first detected SHG signal, the green arrow shows the first fluorescence signal that colocalizes with the SHG signal (see movie 1 series b). Insert: snapshots of series a nucleation point at time 00:00 (hh:mm) (SHG only, red arrow), at time 01:44 where SHG colocalizing fluorescence signal appears (red and green arrow), at time 03:20 showing the fully grown type I.C Xpa crystal (scale bar 10µm). (D) Top row: spinning disk fluorescence based 2D slices snapshots of a type I.C Xpa crystal (see movie 2). The green arrows show the area in the cytoplasm where the crystal appears and grows. the desaturated last frame of the row shows inner and surface defects on the fully grown crystal (scale bar 10µm). Bottom left: segmentation derived type I.C volume change over time, bottom right crystal segmentation of the fully grown Xpa type I.C crystal extracted from movie 3 (scale bar 1 µm).

The type II Xpa crystals are those emerging from multiple simultaneous nucleation sites within the cytoplasmic compartment (Fig. 1C column 2, Fig. 2A and B third row). This morphology was first identified when were observed large and highly fluorescent dense regions of Xpa that produced second-harmonic light with multiple orientations, suggesting crystalline domains. In addition, the rest of the compartment was devoid of soluble Xpa fluorescence, as noticed for type I crystals. These data suggest that multiple nucleation sites could emerge simultaneously and that the subsequent crystal growth could result in a merged polycrystal if these sites were close enough in the intracellular compartment (Supplementary Fig. 3A and Supplementary movie 2). Less frequently, two crystals from more distant nucleation sites within the cytoplasm (for which crystal growth were not interfering with each other) were observed (Supplementary Fig. 3B and Supplementary movie 3, Supplementary Fig. 3C). In figure 2B, the segmentation of a cryoSXT tomogram of two close Xpa crystals (Fig. 2B third row) strongly suggests that two close and independent nucleation sites can mutually impede crystal growth by depleting soluble Xpa between the two crystal interfaces (lower cross section, bottom of the segmentation) as well as resulting in the merging of both crystals (higher cross section, top of the segmentation). The reason for multiple proximate nucleation sites that results in polycrystalline material) in specific regions of some cell cytoplasm remains to be elucidated.

A third Xpa crystalline condensate type, type III, is depicted in figure 1C third column and in figure 2 C and D. From fluorescence microscopy (single or two-photon) images, they can be described as a single region of the cytoplasm in which Xpa is found condensed but without crystal regular flat outer faces. However, type III Xpa condensed phases exhibit multiple SHG signal domains that suggests some localized crystalline ordered regions within these dense condensates (Fig. 1C third column). CryoSXT tomograms, combined with prior cryofluorescence microscopy-based cryoEM grid screening, revealed several carbon dense and highly fluorescent structures that were not previously reported in literature (Fig. 2C right panel). These structures contain two main phases based on the carbon density distribution: one extended, granular or diffuse that engulfs the second phase with a carbon density distribution reminiscent of the large regular (Type I) crystal cores, but with no well-defined planar outer surfaces. A mosaic of four cryoSIM-correlated cryoSXT tomograms of a unique cell (Fig. 2D) allows for the direct comparison between a type I.C Xpa crystal and type IV Xpa crystalline condensate. This condensate exhibits subdomains presenting angular, or partially planar surfaces (one example is marked by an arrow on the segmentation in Fig. 2D), that could result from potential localized nucleation sites within the dense phase. This observation is reminiscent of several ordered domains within the dense Xpa phase detected with SHG microscopy (Fig. 1C column 3). As no dynamical measurements of the stochastic *in vivo* Xpa crystallization process in each cell observed with the cryoSXT technique could be performed prior the EM grid vitrification, no statement can be made on the time point at which these structures have been cryofixed (i.e. as a transient state or a final stable Xpa state), in other words which stages from live fluorescence microscopy observations of this type of ordered Xpa solid condensates (see below, Fig. 3A series c and series d and movie 1) have been trapped during the vitrification process. FRAP experiments performed on these geometrically ill-defined Xpa condensates show no fluorescence recovery as for other types of crystals suggesting that these condensates might be a mixture of crystalline and amorphous dense phases of the protein (Fig. 2E and Supplementary Fig. 3D). For all the type of crystal/crystalline condensates of Xpa, no fluorescence recovery from FRAP experiment has been found as shown in Figure 2F, indicating that the *in vivo* transition of Xpa from the liquid diluted state to the solid condensed state adopt several final morphologies. A better understanding of the multiple pathways leading to all the final condensed stages of Xpa, a more detailed description at various spatial scales has to be performed.

The last observed type of Xpa crystals, type IV, is found in Xpa containing cytoplasmic vesicles (Fig. 1C column 4). This type of Xpa crystals has not been reported before and is described in more details below.

Overall, the morphological description of *in vivo* grown Xpa crystals within HEK293FS cells revealed by our approach is much more elaborated than in previous studies. Whether this is a direct consequence of the use of this specific cell line is, as yet, unclear. However, some of these results are strongly reminiscent of what is regularly observed during *in vitro* crystallization screenings of a purified protein as far as crystals, crystalline aggregates or multiplicity of nucleation sites are concerned. *In cellulo* diffraction experiments were performed on Xpa crystals containing cells and showed that the diffraction properties of Xpa crystals within a cell population span a very wide and continuous range from no diffraction at all to high diffraction quality (data not shown), allowing an *in vivo* grown Xpa crystal structure determination at 1.59 Å resolution (PDB accession number 9GA0), one of the highest structure resolution for an *in vivo* grown crystal protein (Supplementary Fig. 4 and table 1)^2,5^.

### 3 *in vivo* grown Xpa crystals growth exhibits one or two rates of growth and membranes might be involved in the nucleation process

High temporal resolution dynamics of *in vivo* grown crystals are required for the understanding of their ultimate morphology variability as well as for localizing their nucleation sites in the cellular environment. Multiphoton microscopy and spinning disk fluorescence microscopy were used at the single cell level to probe Xpa crystal growth dynamics in living cells. To be able to monitor stochactic crystallization events, a 2 minute time interval resolution was chosen as a minimum temporal unit for sampling a few tens of cells that were chosen for their high Xpa expression levels and their moderate sizes. A typical experiment is depicted in figure 3A. Four Xpa crystallization events out of eighteen chosen cells (each cell being assigned to a series letter) were monitored individually and continuously during 16 hours. In the first cell (series a), crystallization of Xpa into large (15 µm in length) type I.C crystal was observed, while a small (8 µm) type I.C crystal appeared in the cell of series b. The difference observed in the final crystal volume arises from the total amount of soluble Xpa protein available prior the crystallization process that is determined by the volume of the cell and Xpa expression level. The area of the largest ‘series a’ cell cross section at the beginning of the experiment is 535 µm^2^ with a mean fluorescence intensity of 1661 a.u. while these values are 143 µm^2^ and 932 a. u. for the ‘series b’ cell respectively. The growth rate of the series a crystal appeared higher compared to one in series b (Fig. 3A) suggesting that a higher available protein concentration in the cytoplasm would be the main driving force for the speed of crystal growth.

Two additional Xpa crystal growth events, from type III were also observed (series c and series d). Both events originated from the inner plasma membrane vicinity of their respective cells and grew toward the inner part of the cell (Fig. 3A series c and d and movie 1). Interestingly, 40 minutes before crystal growth within series a, the plasma membrane of the cell is locally deformed inwards as shown by the decrease of dilute Xpa fluorescence, in the vicinity of Xpa crystal localization (Fig. 3B and movie 1 panel a). These three observations taken together allows us to hypothesize a potential role of intracellular membranes in Xpa crystal nucleation, maybe acting as an Xpa adsorption interface facilitating the nucleation step^23^ before the crystal tangential growth^24^ . On the contrary, before the type I.C crystal growth in the series b cell, a small (1 to 2 µm sized) SHG punctum was detected at the beginning of the experiment (Fig. 3C). One hour later, the fluorescence signal built up on that puctum, stayed stable for one more hour before undergoing fast crystalline growth while soluble Xpa was simultaneously depleted from the cytoplasmic reservoir. This suggests that an Xpa crystal nucleus might remain stable within the cytoplasm before triggering fast crystal growth from the soluble Xpa reservoir. Such latency is surprising at this stage considering the fast *in vivo* Xpa crystals growth rate and suggests that the nucleus is isolated from the cytoplasmic soluble reservoir before undergoing crystal growth.

The overall dynamics of XPA growth rate can be plotted over time (Fig. 3A right panel) and exhibit two main behaviors: a two-phase (successive fast and slow phases) growth rate for series a (type I.C) and c (type III) and a single fast rate for series b (type I.C) and series d (type III). The fast rate spans from about 70 µm^2^ per minute down to 10µm^2^ per minute. The slow growth rate for both series a and c are about 0.15 µm^2^ per minute. Due to the fact that the nucleation point of series b was spotted, the whole crystal growth rate curve was plotted (Fig. 3C), revealing a sigmoidal shape curve similar to what has been described for *in vitro* crystallization of macromolecules^25–28^.

The overall distribution of crystal growth rates at the cell population level on 32 recorded crystallization events showed that about 62% of these crystals took less than 15 minutes to reach their maximum sizes, when 25% took between 15 and 30 minutes and 13% between 30 and 45 minutes (Supplementary Fig. 5A). Similarly to what has been observed previously^9^ , the crystal formation is concomitant to all available soluble Xpa depletion from the crystallization compartment (Supplementary Fig. 5B).

Such a live crystal growth event was live recorded in three dimensions using spinning disk fluorescence microscopy (Fig. 3D and movies 2 and 3) with a similar time increment of slightly more than 2 minutes. This crystal adopted type I.C geometry and presented inner defects that are characteristic of its class (Fig. 3D desaturated spinning disk fluorescence microscopy image and segmentation). Unfortunately, only part of the crystal was recorded during its growth, the nucleation point being located below the first plane of the acquired volume. Volume segmentation of the visible part of the crystal, plotted over time, showed a fast increase of the crystal growth during the first 5 minutes suggesting that the concentration of the cytoplasmic soluble pool of Xpa was high enough to catch the end of the fast phase. The fast crystal growth rate was measured at 40µm^3^ per minute (Supplementary Fig. 5B) within the range of area growth rates measured by SHG microscopy. A slow growth rate was then observed, probably due to the depletion of the residual soluble pool of Xpa in the reservoir, slowing down the crystal growth until crystallization rate decreased to zero. This measurement allowed us to trap the final phase of the sigmoidal Xpa crystal growth rate and supports what was measured with the single slice-based multiphoton microscopy.

### 4 Xpa is engulfed within vesicles and can crystallize inside them

Recently, fluorescent puncta have been observed within cells of transgenic C elegans^10^ . Interestingly, using fluorescence confocal microscopy, we also detected numerous Xpa dense puncta within both highly Xpa and poorly Xpa expressing cells (Fig4A and Fig. 4B). Labeling experiments with ER tracker showed that these Xpa dense puncta were surrounded by ER derived membranes. Moreover, concentration of Xpa in these vesicles is higher compared to the concentration of Xpa in the cytoplasmic compartment (Fig. 4A). These Xpa enriched vesicles are also observed even when the cytoplasm is devoid of soluble fluorescent protein (Fig. 4B) suggesting that this could be an early cell response to heterologous Xpa expression. The typical diameter of these vesicles is in the range of 1 µm but can reach several microns (see below Fig 4D). This observation suggests that Xpa is stored inside vesicles upon expression via an unidentified mechanism. This phenomenon might contribute to the lack of toxicity that could arise from the overexpression of the fluorescent protein in *C elegans* as noticed by Kuramochi et al.^10^. Within such a confined environment, Xpa is expected to be highly enriched and concentrated which could turn out to be highly favorable for Xpa crystallization (similarly to the conditions required for *in vitro* crystallization trials). Indeed, Xpa was found to be able to form crystals in these vesicles as shown in Fig. 4C where the relative fluorescence level in all these vesicles across the cell are similar with one exception for which the fluorescence intensity was more than four times higher (in agreement with Supplementary Fig. 1D). Similar observations of µm-sized SHG and fluorescent positive puncta were made with SHG microscopy (Supplementary Fig. 7A and Supplementary movies 3 and 4). In addition, using live multiphoton microscopy, a 6µm-sized homogeneous rhombic shaped crystal was monitored over time and found to be embedded within a vesicle (Fig. 4D left panel and movie 4) while a homogeneous signal of Xpa was still observed throughout the cell, suggesting that a physical barrier isolated the crystal from the cytoplasm. A similarly sized crystal engulfed in a vesicle was also observed on cryoSXT tomograms (Fig. 4D right panel). This vesicle lies next to a large type I.C crystal that collected all the cytoplasmic pool of Xpa as seen by cryo SIM fluorescence microscopy (Supplementary Fig. 7B) indicating that this small Xpa crystal was only derived from the protein pool included in the vesicle. CryoSXT tomography also revealed two types of flat shaped objects inside vesicles. The first category, depicted in Fig. 4E, can be described as flat shaped plates surrounded by vesicle membranes. They are found in a small subset of cells (10 over almost 150 recorded tomograms), and some of them are connected to a diffuse carbon dense structure within the vesicle (Fig. 4E lower panel). The second category consists of needle like objects also surrounded by vesicle membranes (Fig. 4F). These structures are reminiscent of *in vivo* grown luciferase crystals previously described in insect cells which could reach 200 µm in length ^29,30^ which could span the whole cell extending their plasma membrane. A similar pattern could be seen in Fig. 4F (upper part), where the vesicle membrane is distorted from its regular circular shape to an oval extended shape. Despite our efforts, because of the small size of these two types of structure (plates and needles) and the patterning effect of cryoSIM data that appeared on the global fluorescent background due to cytoplasmic pool of Xpa, an unambiguous correlation with Xpa fluorescence was not possible. However, the size of these objects within vesicles was comparable to the SHG signal-and fluorescence positive puncta observed with multiphoton microscopy (Fig. 4G and movie 5, Supplementary Fig. 7A). These objects were often seen gathered in specific places in the cytoplasm or sometimes observed isolated. FRAP experiments performed on part of these small puncta showed no fluorescence recovery revealing their non-diffusive solid nature (Supplementary Fig. 7C). All these observations suggest that Xpa can be engulfed inside vesicle within the cytoplasm and can crystallize in these vesicles into different crystalline morphologies. These crystals in vesicles have been classified as type IV.

**Figure 4:**
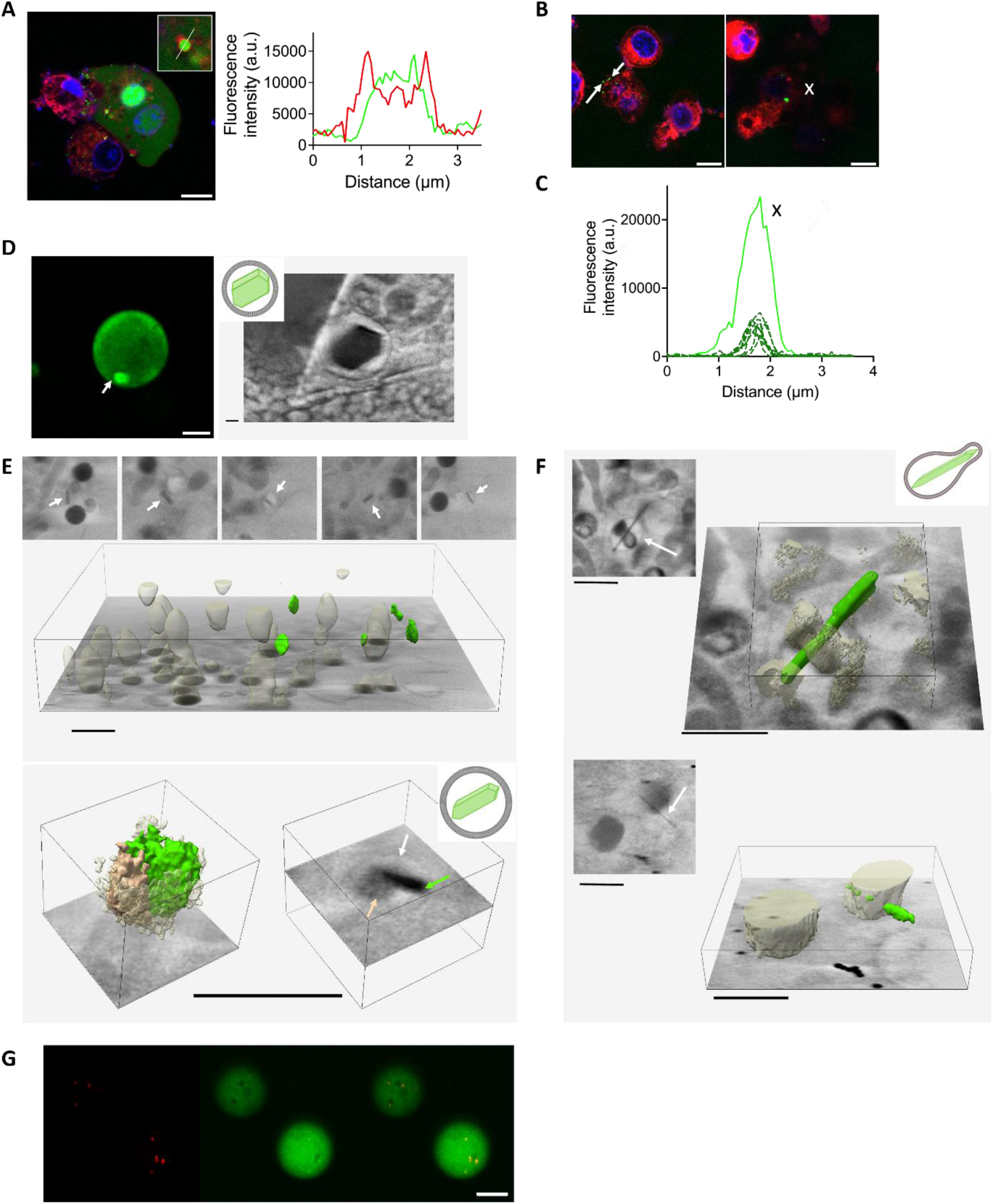
Type IV Xpa crystals. **(A)** Confocal fluorescence 2D slice of a highly Xpa expressing cell. Green: Xpa fluorescence channel, red: ER tracker fluorescence channel, blue: Hoechst fluorescence channel (scale bar 10µm). Insert: Xpa enriched ER membrane derived vesicle. Plot profile of the intensity of the red and green channels along the line shown in the insert. (B) Confocal fluorescence 2D slices of a low Xpa expressing cell (scale bar 10µm). White arrows show two Xpa enriched vesicles and the white cross show a highly Xpa fluorescent vesicle. (C) intensity of Xpa fluorescence within vesicles, vesicles with moderate Xpa fluorescence are in dashed line and dark green, highly Xpa fluorescent vesicle marked with a cross on the 2D plane in continuous light green line. (D) Left: multiphton fluorescence snapshot of a type IV Xpa crystal (see movie 4). The arrow shows the Xpa fluorescence depleted inner part of the vesicle (scale bar 10 µm). Right : area of a slice of a 3D reconstructed cryoSXT tomogram showing a Xpa crystal within a vesicle (see supp Fig. 7B) (scale bar 1µm). (E) Top: areas of 2D slices from a reconstructed cryoSXT tomogram showing plates engulfed within cytoplasmic vesicles. The white arrows show the vesicles membranes. Middle segmentation of the 3D reconstructed cryoSXT tomogram.semi transparent beige structures are lipid droplets and green structures are carbon dense plates within vesicles shown in top (scale bar 1µm). Bottom : 3D segmentation of a carbon dense plate found in on vesicle from the tomogram in the middle panel. Green structure : carbon dense plate, beige opaque structure : intravesicular carbon dense region next to the plate, semi transparent white structure: vesicle membrane. On the left a 2D slice of the 3D segmentation, the arrows show the carbon density segmented in the left panel with the same color code (scalebar 1µm). (F) two examples of needles found in two 3D reconstructed cryoSXT tomograms. Areas of 2D slices and the corresponding segmentation are shown. The white arrows show carbon density of needle containing vesicle membranes, green segmentated structures are the needles, semi transparent brown segmented structures are carbon dense structure found in the subvolume. (G) Movie 5 snapshot of SHG, multiphoton fluorescence, and merged channels showing the colocalization of Shag and Xpa fluorescent punctae in two cells. (scale bar 10µm).

### 5 Xpa crystallization occurs also within Xpa enriched externalized vesicles

As seen previously, cytoplasmic Xpa is sequestered at high concentration inside vesicles. We noticed that such vesicles could be expelled from the cell. Indeed, using confocal microscopy, accumulation of Xpa enriched vesicles is often observed concentrated on the periphery of the cell, and also some of them appeared to be expelled from it (Fig. 5A). In the case of highly expressed cytoplasmic Xpa (i.e. homogeneous Xpa distribution throughout the cytoplasm), large vesicles can bud from the plasma membrane and might be released in the extracellular medium (Fig. 5A down left and middle panels). In cryoSXT tomograms, some Xpa crystals surrounded by membrane were observed outside of a cell, directly lying on the carbon layer of the cryoEM grid (Fig. 5B and C). Oppositely to intracytoplasmic grown crystals, these are devoid of internal inclusions coming from the crowded intracellular environment suggesting that the crystallization process occurred directly within the externalized vesicle. To support this hypothesis, fluorescent puncta were observed inside such vesicles outside of the cells (Fig. 5A right panel and movie 5) which could be premises of Xpa crystal nuclei. An interesting observation of such an externalized crystal (Fig. 5B) suggests that these extracellular vesicles could be fused to other expelled carbon rich vesicles and then potentially provide additional material for further Xpa crystal growth. In figure 5C, multiple Xpa crystals are seen inside an externalized vesicle, probably originating from distinct Xpa crystal nuclei. Interestingly two of these crystals exhibit curved surfaces that follow the shape of the membrane, reminiscent of that which was observed in figure 3A for both Type III crystals (series c and d) that nucleate in the vicinity of the plasma membrane.

**Figure 5:**
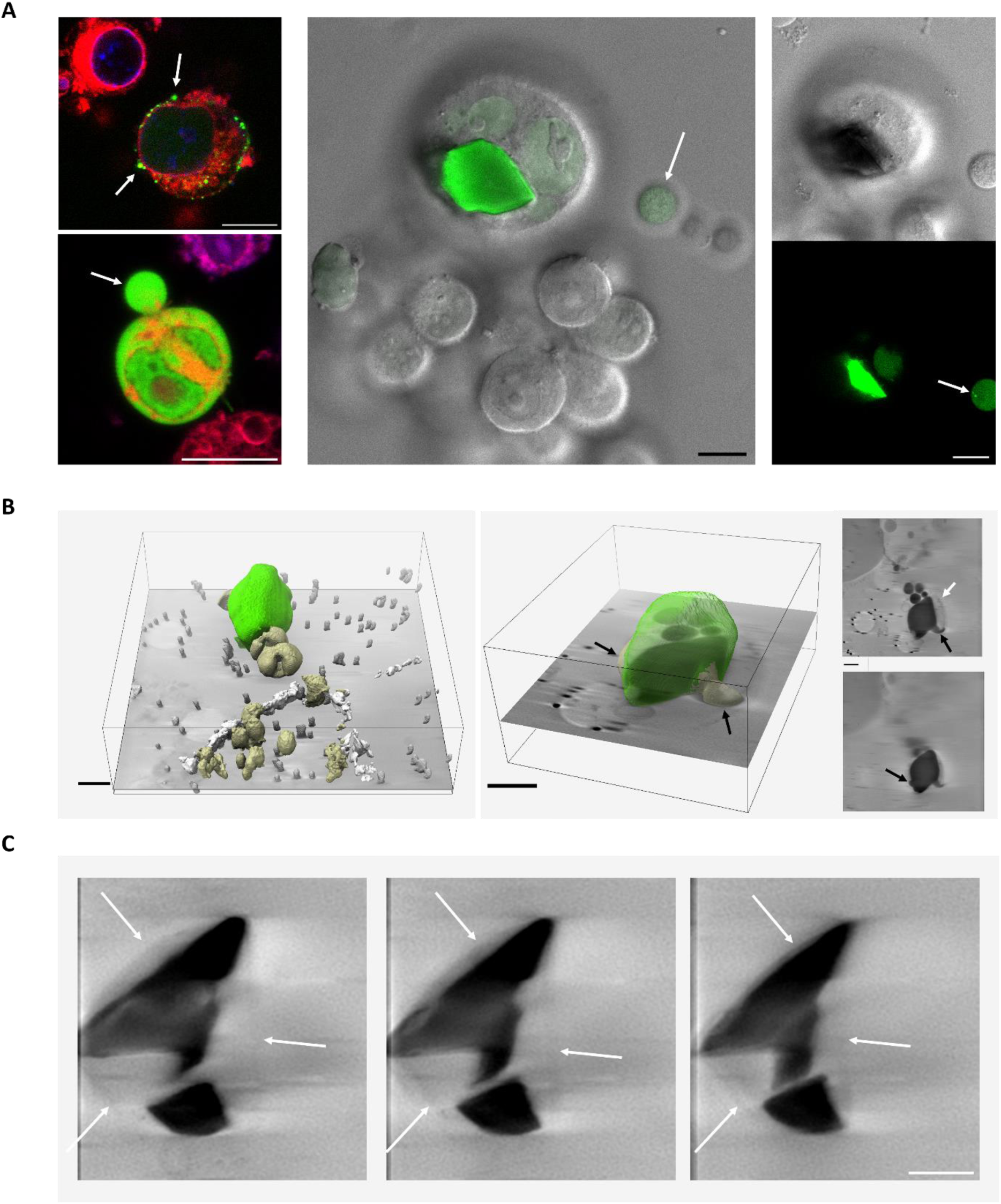
Xpa crystallization in extracellular vesicles. (A) left : confocal fluorescence images of a low Xpa expressing cell (top) and a high Xpa expressing cell (bottom). Arrows show Xpa rich vesicles at the plasma membrane of the cells. Green channel : Xpa fluorescence, Red channel: ER tracker fluorescence, blue channel: Hoechst. Middle: merged fluorescence (green) and DIC (grey) channel of a type II Xpa crystal in a cell. The white arrow shows a Xpa enriched extracellular vesicle. Right: DIC (top) and fluorescence (bottom) channels 2D slice of the cells in the middle panel. The white arrow shows fluorescent condensates within the vesicle (see movie 6). All scale bars are 10µm. (B) left: segmentation of a 3D reconstructed cryoSXT tomogram of a Xpa crystal within an extracellular vesicle: green structure: Xpa crystal, white structure: cell membrane, beige structures: carbon dense vesicles, grey structures: gold fiducials. Middle: segmentation os a subvolume from the left panel. Semi transparent green: Xpa crystal, beige structures pointed with black arrows: carbon vense vesicles attached to the crystal. Right panel: two 2D slices from the cryoSXT tomogram showing the vesicle membrane (white arrow) and both carbon dense Xpa crystal attached structures (black arrows). All scale bars are 1µm. (C) 2D slices from a 3D reconstructed cryoSXT tomogram of a type II Xpa crystal within an extracellular vesicle. White arrows indicates the membrane of the vesicle. Scale bar 1µm.

### 6 Fast decrease of cell volume induces plate and needle shaped Xpa crystallization as well as dendrite shaped Xpa aggregation

The global cell volume was observed to decrease by roughly 30% over one to three days after transfection (Fig. 6A). This phenomenon suggests that the concentration of Xpa, reached one day post transfection within the cytoplasm, increases while the cell volume decreases and that conditions for Xpa crystallization would become more favorable with time. A fast decrease of the cell volume can be achieved by establishing a hypotonic stress by adding sorbitol in the medium, as shown by Boyd-Shiwarsky et al.^31^. To reach osmotic equilibrium, water molecules are then expelled from the cell towards the culture medium, rapidly reducing their volume. In our hands, the addition of 0.5M sorbitol in the cell culture medium induced a cell volume shrinkage of about 50% within 30 minutes (Fig. 6B). Upon cell shrinkage Xpa underwent a phase change, as seen by fluorescence microscopy, by condensing Xpa into plate- and needle-like structures, and, less frequently, into dendrite-like structures. Plates could be found alone (Supplementary Fig. 8) of mixed with needles (Fig. 6C top panel). As opposed to what was observed for Xpa crystal formation without sorbitol, Xpa plates and needles coexisted with the soluble pool of Xpa in the cytoplasm (Fig. 6C top panel). In addition, dendrite structures, that could also be found together with plates (Fig. 6C down panel), concomitantly depleting all the remaining pool of soluble Xpa in the cytoplasm (movie 6). Surprisingly, whenever the dendritic phase was formed, dissolution of the needles and, to a lesser extent, the plates was observed (movie 6). It is noteworthy that dendrite structures largely extend along the plasma membrane (movie 6). SHG microscopy showed that Xpa (within the plates, the needles as well as the dendrite structures) was ordered (Fig. 6D) and no fluorescence recovery from FRAP experiment performed on these three types of structures was observed (Fig. 6E), indicating their solid nature. However, no X-ray diffraction pattern was observed in Xpa containing cells submitted with sorbitol, suggesting a poor crystalline intrinsic order at the molecular level (data not shown).

**Figure 6.**
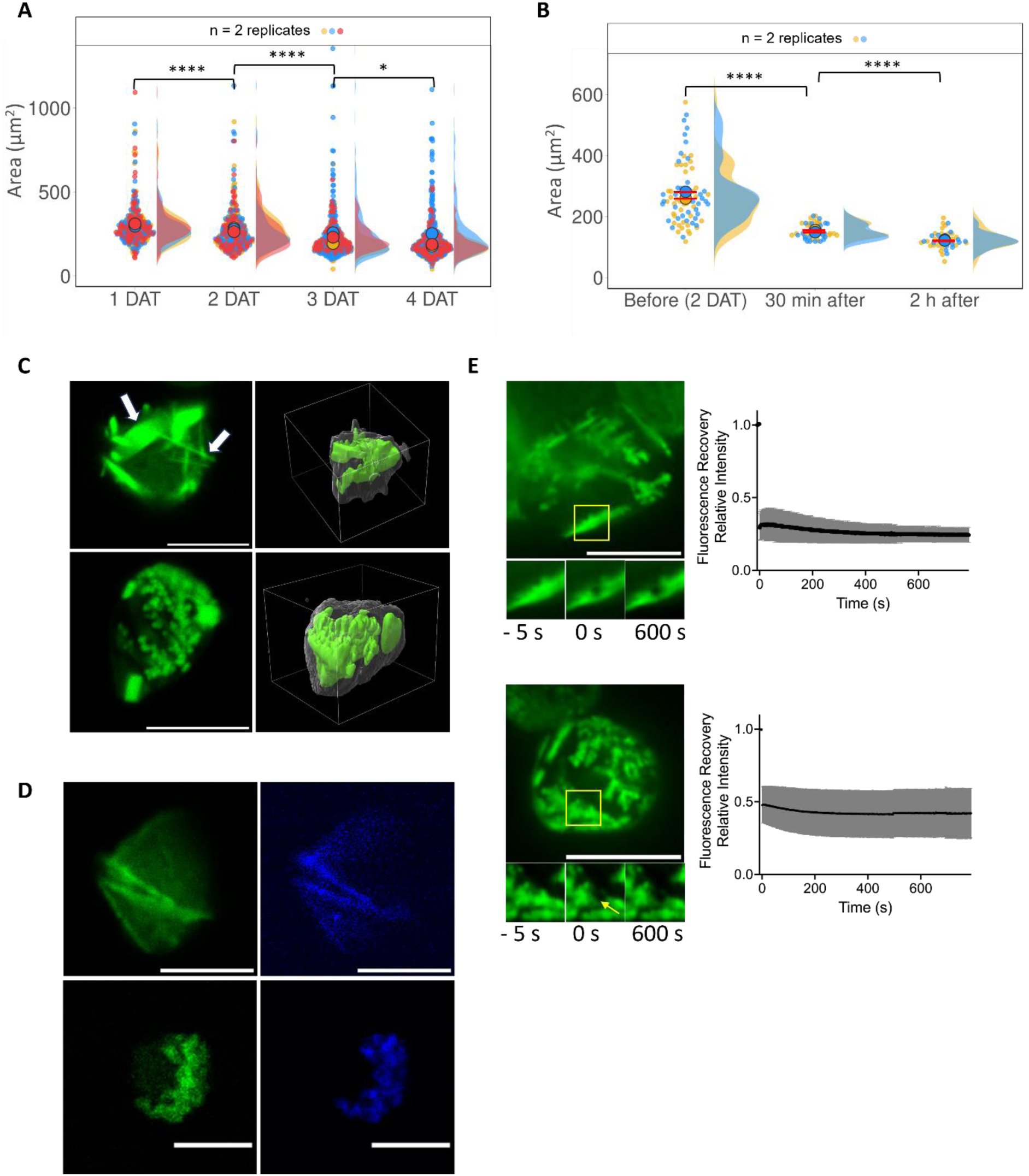
Effect of fast cell volume decrease on Xpa crystallization and aggregation. (A) Quantification of cell size as a function of day post-transfection measured in images recorded using multiphoton fluorescence. Three replicates; p-values between 1 DAT and 2 DAT, 3 DAT are lower than 0.0001, whereas those of between 3 DAT and 4 DAT is 0.0135. (B) Quantification of cell size before and after addition of sorbitol measured in images obtained using confocal fluorescence microscopy. Two replicates; p-values are lower than 0.0001. (C) Confocal fluorescence image (left panel) and their segmented volume (right panel) of Xpa plats (top) and Xpa dendides (bottom) formed after addition of sorbitol. Scale bar 10 μm. (D) Multiphoton fluorescence image (left panel) and SHG image (right panel) of Xpa plating (top) and Xpa dendides (bottom) formed after addition of sorbitol. Scale bar 10 μm. (E) Fluorescence images with plates (upper line) and dendrites (lower line) of a representative cell analyzed by FRAP. Scale bar: 10 μm. Quantification of the FRAP data from 5 for plates and 15 for dendrites different experiments.

## Discussion

We have demonstrated that expression of Xpa within the HEK293FS cell line is an excellent model to get more insight on *in vivo* macromolecular crystallogenesis. Combined with a multitechnical/ multimodal approach, an atlas of 4 types of *in cellulo* grown crystals has been established. A recent paper has also suggested that Xpa crystals are not only present as types I.C and I.N but can also form crystalline puncta ^10^. In our experiments, such puncta are surrounded by ER derived vesicular membranes and some of them are SHG positive indicating an intrinsic molecular order. Unexpectedly, low expression levels of Xpa were sufficient to elicit the compartmentalization of the protein into these vesicles. We did not investigate the early cell response pathway to Xpa expression but Tsutsui et al. have shown that cytoplasmic *in vivo* grown Xpa crystals could be eventually surrounded by autophagic membranes and sometimes expelled from the cell ^9^. An autophagic response to Xpa expression could also be the cause of the protein engulfment within the vesicles observed in our experiments. This might also be a reason for the non-toxic effect of Xpa expression within c. elegans^10^. This compartmentalization of Xpa might explain why the SHG/fluorescent positive puncta do not gather all the cytoplasmic soluble pool of Xpa into large crystals despite the rapid kinetics of crystal growth due to the physical barrier between the intravesicular crystalline Xpa and the cytoplasmic reservoir.

Two types of small intravesicular structures observed on cryoSXT tomograms could be good candidates for small Xpa crystals, namely plate-like structure or needle like structures. Even though no direct correlation with cryo fluorescence microscopy could have been made in our measurements, Xpa is able to crystallize into similar structures when the cell volume is rapidly decreased upon osmotic shock. Noteworthy, when the vesicle is larger (around 5µm in diameter), intravesicular type I.N like rhombic crystal of Xpa can be observed (Fig. 4C and Supplementary Fig. 7B) that constitutes the third intravesicular crystal form. This suggests that the Xpa crystal geometry might depend on the quantity of available Xpa within a defined volume. It is also worth noting that, upon Xpa precipitation into the dendritic form, preexisting Xpa needles dissolve rapidly to enrich the precipitated phase and, to a lesser extent, the preexisting Xpa plates. This is not surprising considering that Xpa crystals, like most of *in vivo* grown protein crystals^2^, dissolve rapidly when the cell membrane is disrupted and thus are highly sensitive to the surrounding chemical conditions.

We spotted one example where long living SHG/fluorescent-positive punctum seemed to nucleate to form a type I.C crystal (Fig. 3C and movie 1 series b). At first, this punctum could be assigned to a type IV crystal isolated from the cytoplasmic reservoir of soluble Xpa. But the nucleation point of a type I.C crystal requires the intravesicular ordered crystal to disrupt the vesicle membrane and seed the large type I.C crystal. The CryoSXT observation of needles distorting the vesicle membrane from its native spherical shape (Fig. 4E) supports the hypothesis of a needle-like type IV Xpa crystal that grew rapidly, overcoming the membrane rupture point and then reaching the cytoplasmic Xpa reservoir. Such a vesicle membrane would not have the ability to extend in response to such physical constraints contrary to the plasma membrane, as previously demonstrated^29,30^.

A significant number of observations in this work point towards the role of cellular membranes in Xpa crystal nucleation. First, almost all Xpa crystal types were found in contact with membrane structures, the vast majority of type I.C, type II and type III are found connected to plasma membrane or inner vesicle membrane leaflet. Secondly, there are plenty of membrane delimited organelles in the cytoplasm that would account for the far more frequent occurrence of Xpa crystallization in the cytoplasm compared to the nucleus where only the inner nuclear membrane might be involved in Xpa crystal nucleation. Thirdly, for crystals that seem not in contact with these membranes, they are found embedding membrane delimited organelles and these can be potential nucleation sites. Only high temporal and spatial resolution studies could unambiguously assign the role of membrane structures (surrounding or engulfed within these large crystals) for the nucleation step in this membrane rich environment. We can nevertheless note that in movie 4 (series c and d), small-sized type III crystals originate from the inner plasma membrane region and grow towards the center of the cell. Similarly, all type I.N crystals observed by 3D imaging technique (cryoSXT or confocal microscopy) are in contact with the inner nuclear membrane. Several extracellular Xpa crystal grown inside vesicles showed curved faces that follow the inner vesicle membrane instead of distorting them (Fig. 5C). All the Xpa plates and needles arising upon cell volume shrinkage following sorbitol treatment present contact points with the plasma membrane, whereas the dendrite-like Xpa aggregates follow the inner plasma membrane curvature. Despite all these converging observations, a definitive proof of the role of cellular membranes in Xpa nucleation would require higher resolution techniques such as cryo electron tomography (cryoET) on the very early steps of Xpa nucleation sites.

It would be interesting to apply the approach we presented in this work to other *in vivo* grown crystals, not only in the case of heterologous expression systems, but also in naturally occurring *in vivo* grown crystals to highlight the similarities and differences between all *in vivo* crystallizable proteins and how the cellular environment modulates this process.

At the mesoscale, cryoET is a very powerful method to probe at high resolution the close vicinity of *in vivo* grown crystals and observe crystalline defects ^6^ when such events are frequent in the cell population. Including the cryoSXT in the pipeline for *in vivo* crystallization studies turned out to be essential in our model system. Indeed, it offers the possibility to rapidly acquire entire cells without the need of the cumbersome lamellae preparation required for cryoET. Getting statistics on a whole cell population and spotting easily rare or transient events allowed us to enrich the atlas and describe, at high spatial resolution, the observations using fluorescence-based techniques. Combined with live imaging approaches, the potential of this X-ray absorption imaging technique appeared invaluable in this work. The next step will be to systematically combine cryoSXT with electron tomography while keeping the chemical information from cryo fluorescence microscopy^16,20,22,32^. The *in vivo* grown Xpa crystal in HEK293FS cell line model allows us to set such workflow by selecting the rare events of interest from the hundreds recorded cryoSXT tomograms and mill a limited number of lamellae of the very localized region where the process need to be described at higher resolution via cryoET. For example, scanning the type III crystalline region and get information on the local Xpa molecular lattices to correlate with embedded cellular structures could reveal the role of the cytoplasmic heterogeneity in Xpa crystal nucleation within these Xpa dense phases.

Now that the *in vivo* grown Xpa crystal atlas in its wide diversity has been set and the multitechnical/ multimodal approach for this system has been demonstrated, we are focusing on the very early events of crystal nucleation within the cellular environment. More specifically on the states of the protein that preclude the first crystal nuclei formation and on the suggested role of biological membrane in the nucleation process (manuscript in preparation). Specific local intracellular conditions at the mesoscale that will trigger the formation of all type of Xpa crystals described in this work will also be addressed.

As a final note, the understanding and evaluation of *in vivo* crystallogenesis through pipelines such as the one presented in this work, associated with *in cellulo* diffraction techniques and the recent advent of machine learning based design of crystallizable proteins ^33^ would significantly contribute to this new field of macromolecular X-ray diffraction crystallography while avoiding the bottleneck of classic *in vitro* crystallography.

## Supporting information

Multiple Xpa In vivo crystallization routes in HEK 293 human cells supplementary

## Contributions

F.J and L.C. conceived the project, O.L., C.B. and A.S. performed the sample preparation and characterization. A.S. performed FRAP and spining disk experiments and data analysis. O.L. and A.S. performed multiphoton, SHG and confocal experiments and data analysis; O.L and F.J. and E.P performed cyoSXT experiment. H.C. and O.L. did cryoSXT and fluorescence microscopy segmentation. T.I. carried out Xpa crystal frozen preparation, X-ray data collection and data processing and T.I. built the structure model and refined the structure; S.J. designed FRAP and spinning disk experiments and data analysis. O.L., A.S., E.P., M.P., H.C., F.L., contributed to the discussions and comments on the results; O.L., M.P. and F.J. jointly wrote the manuscript.

## Notes

### Competing Interest Statement

The authors have declared no competing interest.

