## Supplementary material for "Multiple Xpa In vivo crystallization routes in HEK 293 human cells": Multiple Xpa In vivo crystallization routes in HEK 293 human cells supplementary

Supplementary Figures

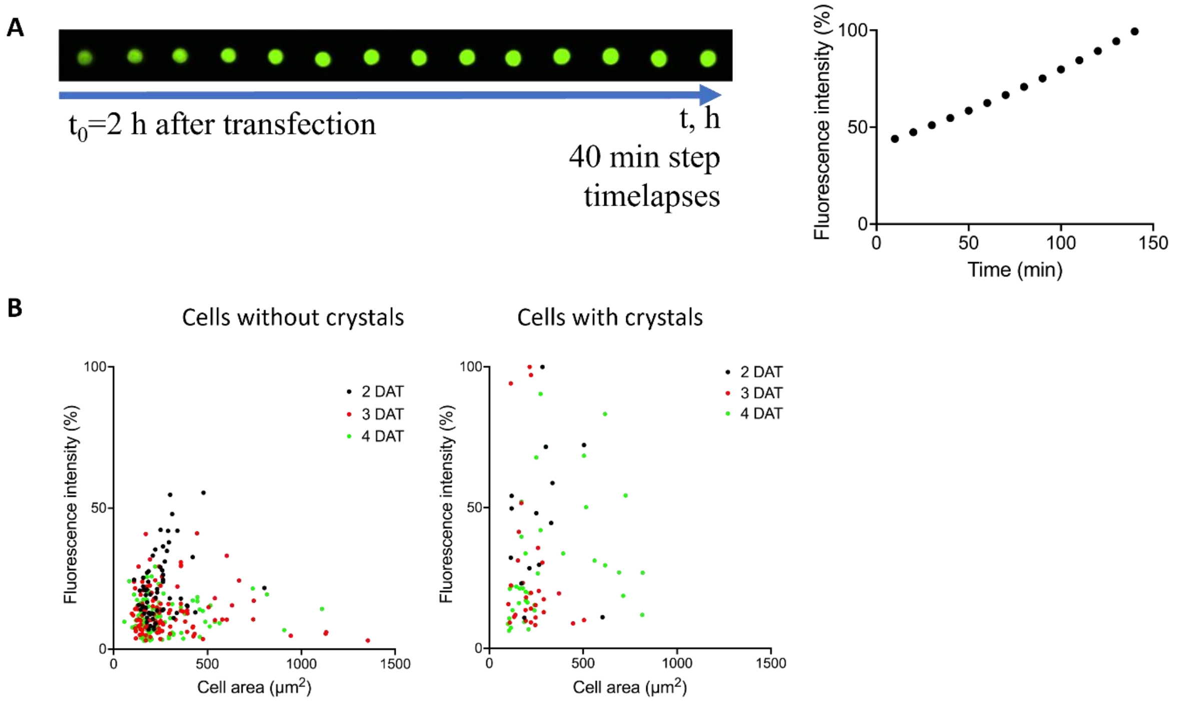

C

| | 2 DAT | 3 DAT | 4 DAT |
|---|---|---|---|
| Amount of crystals, % | 1.2 ± 0.2 | 4.6 ± 0.4 | 17.9 ± 1.7 |

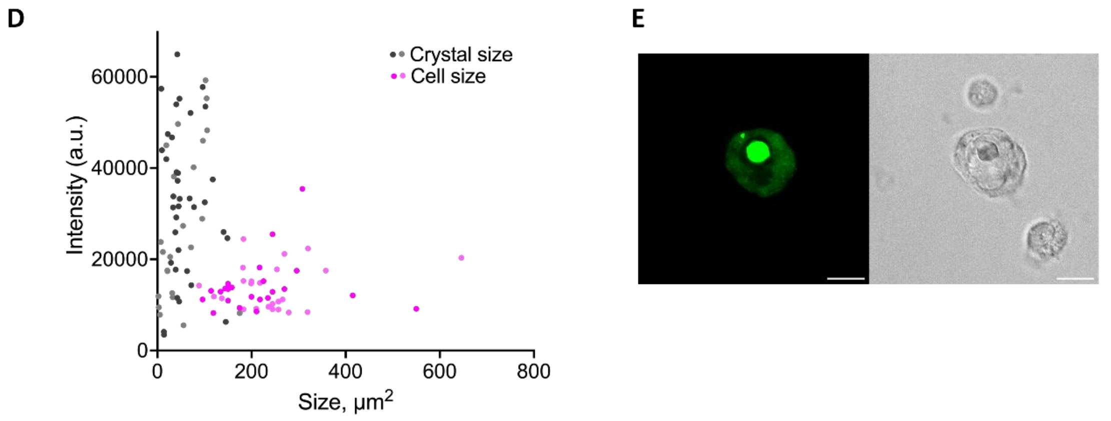

**Supp figure 1. XPA crystallization. (A)** Representative fluorescence images obtained by multiphoton microscopy 3 hours after transfection for XPA. Quantification of the increase in fluorescence signal level for the images on the left. (B) Correlation between the fluorescent signal obtained by multiphoton microscopy and the size of cells without crystals (left panel) and with crystals (right panel), obtained for 2, 3 and 4 days after transfection. (C) Quantification of the increase in the number of crystals in the cell population on days 2, 3 and 4 after transfection. Three replicates, 500 cells each. (D) Correlation between the fluorescent signal obtained by confocal microscopy and the crystal size (gray dots) and cell size (magente dots). 4 DAT, two replicates. (E) Confocal (left) and DIC (right) microscopy images of the I.N type crystal in Figure 1.C showing Xpa fluorescence in the cytoplasm.

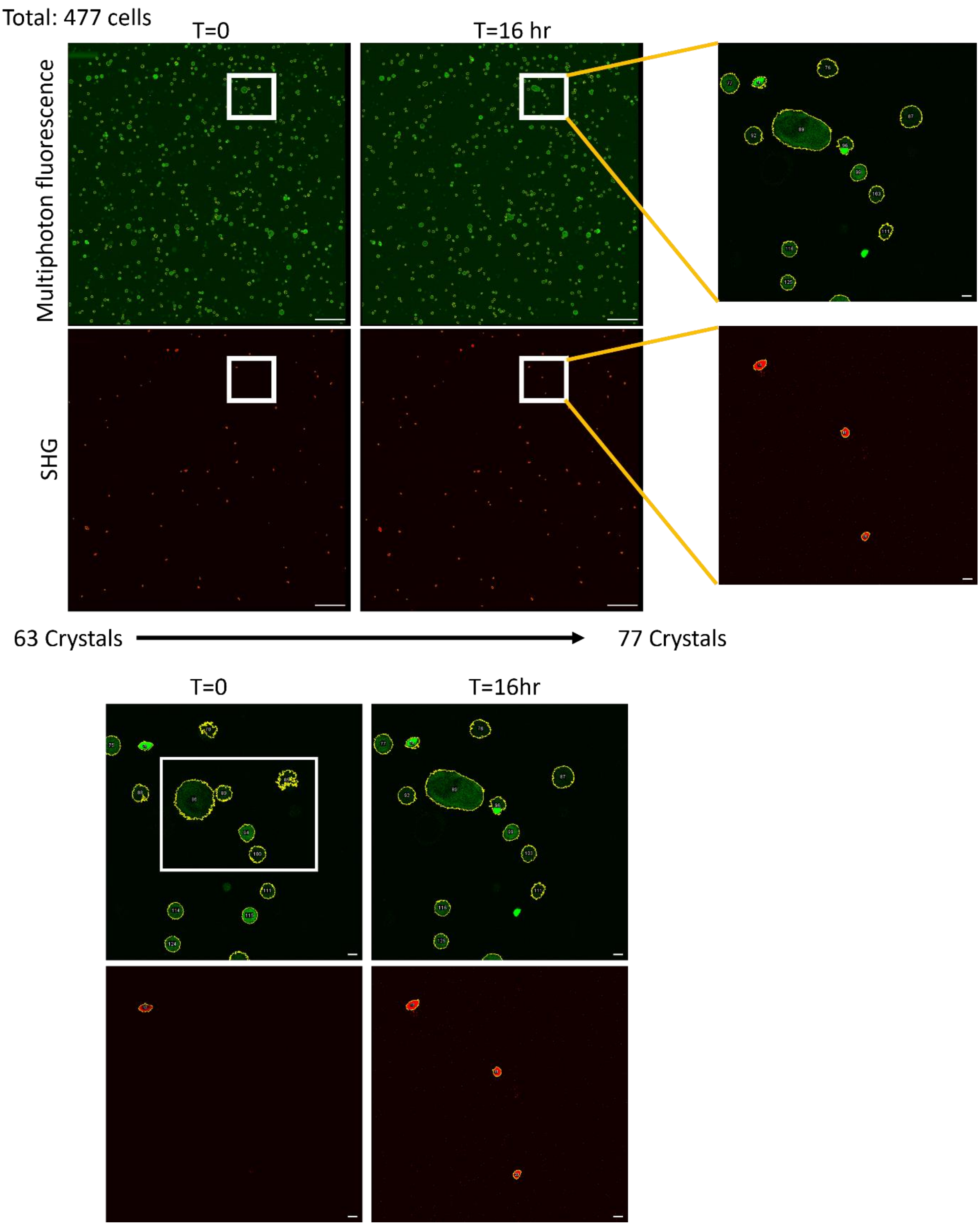

Supp figure 2: Multiphoton based long dynamics of Xpa crystal appearance at the cell population level. Top left: fluorescence (top) and SHG (bottom) channels images of 477 Xpa expressing cells at time 0 and time 16hrs and the increase of the number of cell crystals from 63 crystals at time 0 to 77 crystals at time 16 hrs. Imaging timestep is 18 minutes. (scalebar 200µm). Right: magnified area of left panel at time 16hrs delineated with a white square. Yellow outlines and numbers assigned to fluorescent or SHG structures are the results from the object detection protocol used in Fiji software (see material and methods). Scale bars 10 µm. Bottom: enlarged area from the top images at time 0 and time 16hrs. The white rectangle represent the area used for Supp movie 1. scale bars 10µm.

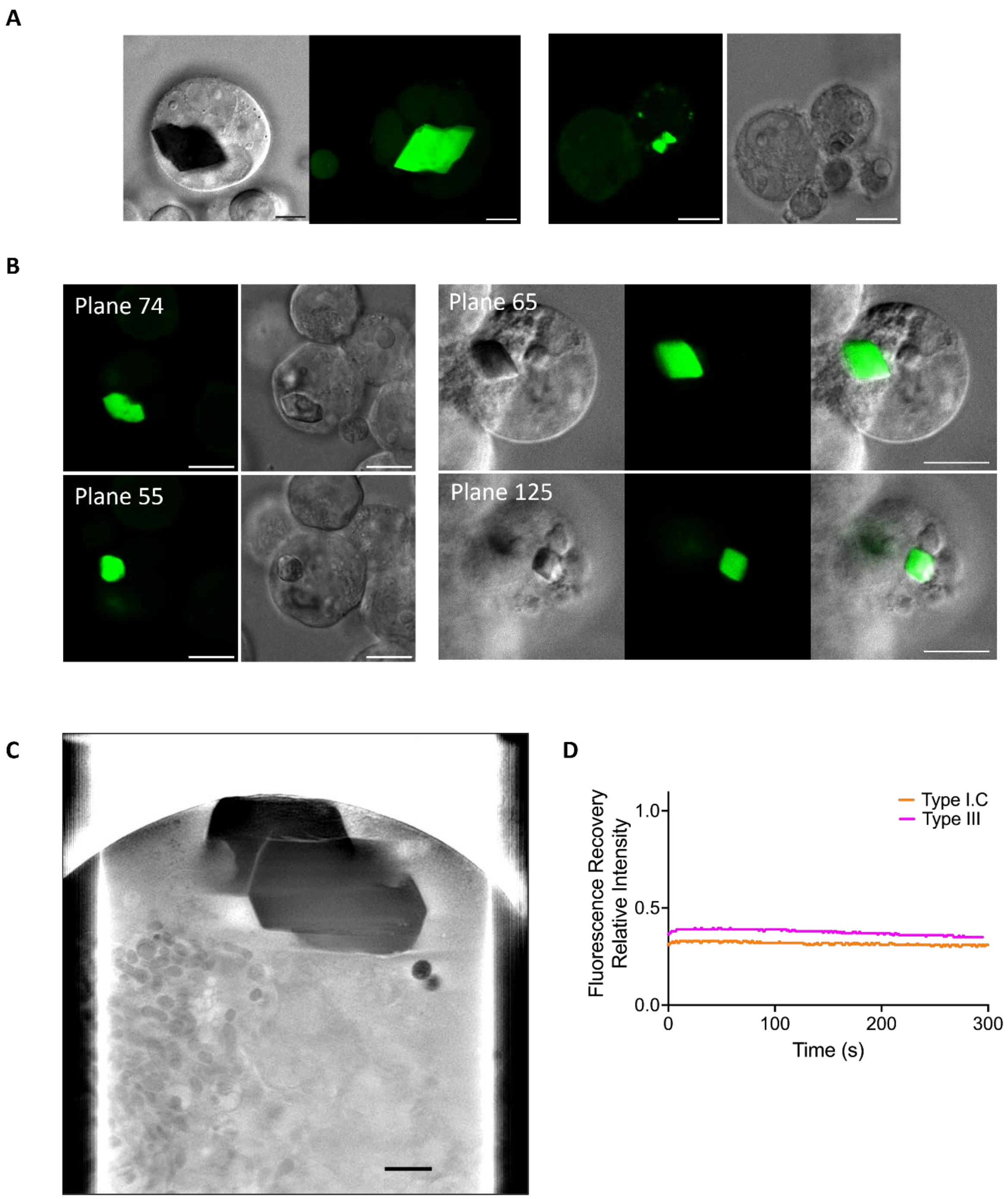

Supp Figure 3: (A) confocal DIC and fluorescence planes of type II Xpa crystals within cells. Left: fused Xpa crystals (see Supp movie 2), right 2 Xpa crystals in contact. (B) confocal DIC and fluorescence different planes of the same cell. Right: Type II (plane 74) and type III (plane 55) Xpa crystals, left: two type I.C crystal (see Supp movie 3), the thirs image is the merge of DIC and fluorescence channel. All scale bars 10µm. (C) 2D plane of a reconstructed cryoSXT tomogram of type II crystal in a cell (see supp movie 4). scale bar 1µm. (D) Quantification of FRAP data for the data in Figure 2(E).

**Table 1 Data collection and refinement statistics (molecular replacement)**

| | PDB ID 9GA0 |
|---|---|
| **Data collection** | |
| Space group | P21212 |
| Cell dimensions | |
| *a*, *b*, *c* (Å) | 50.00, 84.32, 118.35 |
| $\alpha$, $\beta$, $\gamma$ (°) | 90,0, 90.0, 90.0 |
| Resolution (Å) | 19.858-1.61 |
| | (1.78-1.61) * |
| $R_{sym}$ or $R_{merge}$ | 0.26 (1.8) |
| $I / \sigma I$ | 1.2 |
| Completeness (%) | 94.2 (67.5) |
| Redundancy | 13.7 (14.5) |
| **Refinement** | |
| Resolution (Å) | 1.59 |
| No. reflections | 45140 |
| $R_{work}$ / $R_{free}$ | 0.1989/0.2350 |
| No. atoms | |
| Protein | 3566 |
| Ligand/ion | 49 |
| Water | 475 |
| *B*-factors | |
| | 17.96 |
| R.m.s. deviations | |
| Bond lengths (Å) | 0.010 |
| Bond angles (°) | 1.04 |

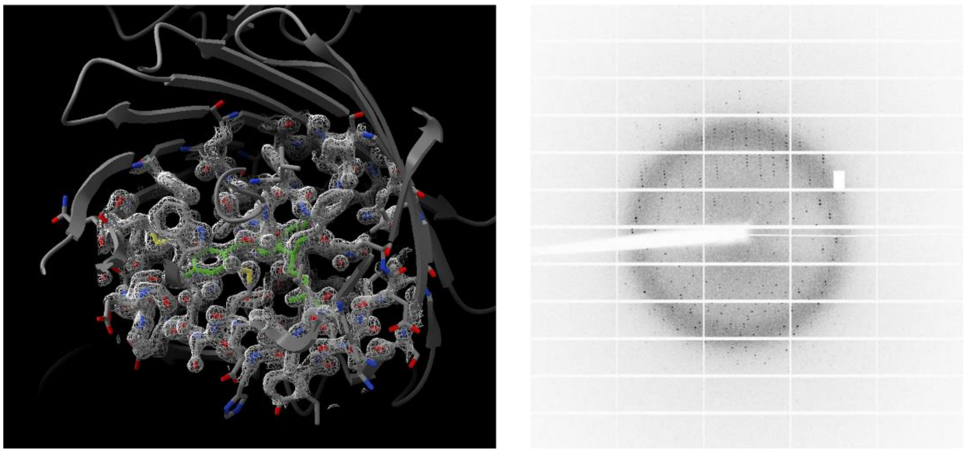

**Supp figure 4 : High resolution in vivo grown Xpa crystal**. Table 1 : X-ray data collection and refinement statistics.

2Fo-Fc electron density map of 9GA0 structure (contour 1.0 s)

Corresponding diffraction pattern recorded on an EIGERX 16M detector on the PROXIMA-1 beamline at Synchrotron SOLEIL.

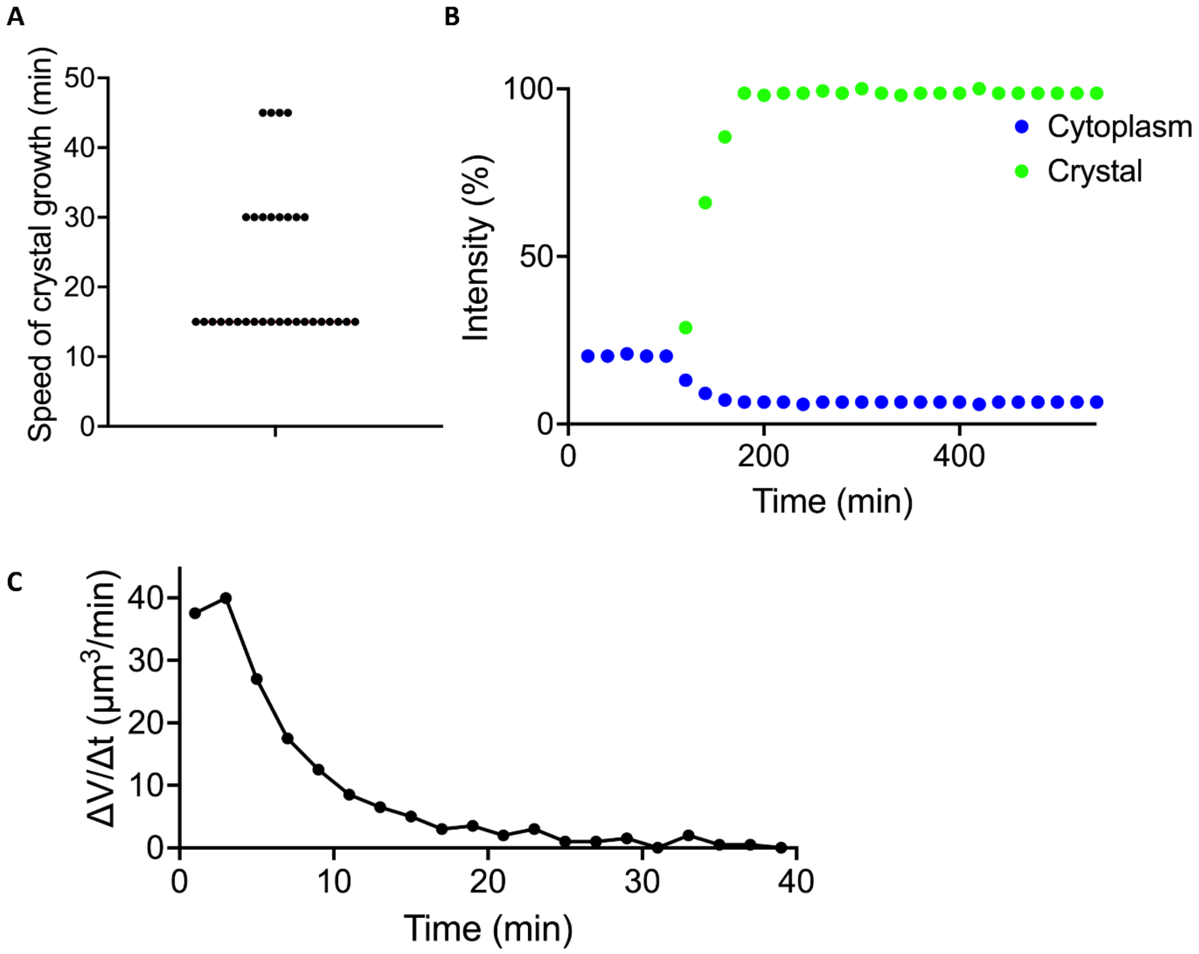

Supp figure 5 : (A) Quantification of Xpa crystal growth rate data. Time step 15 min. (B) Curves of changes in the fluorescence level in the cytoplasm and in the Xpa crystal as the crystal grows. (C) : Type I.C growth rate over time from segmented spinning disk fluorescence data.

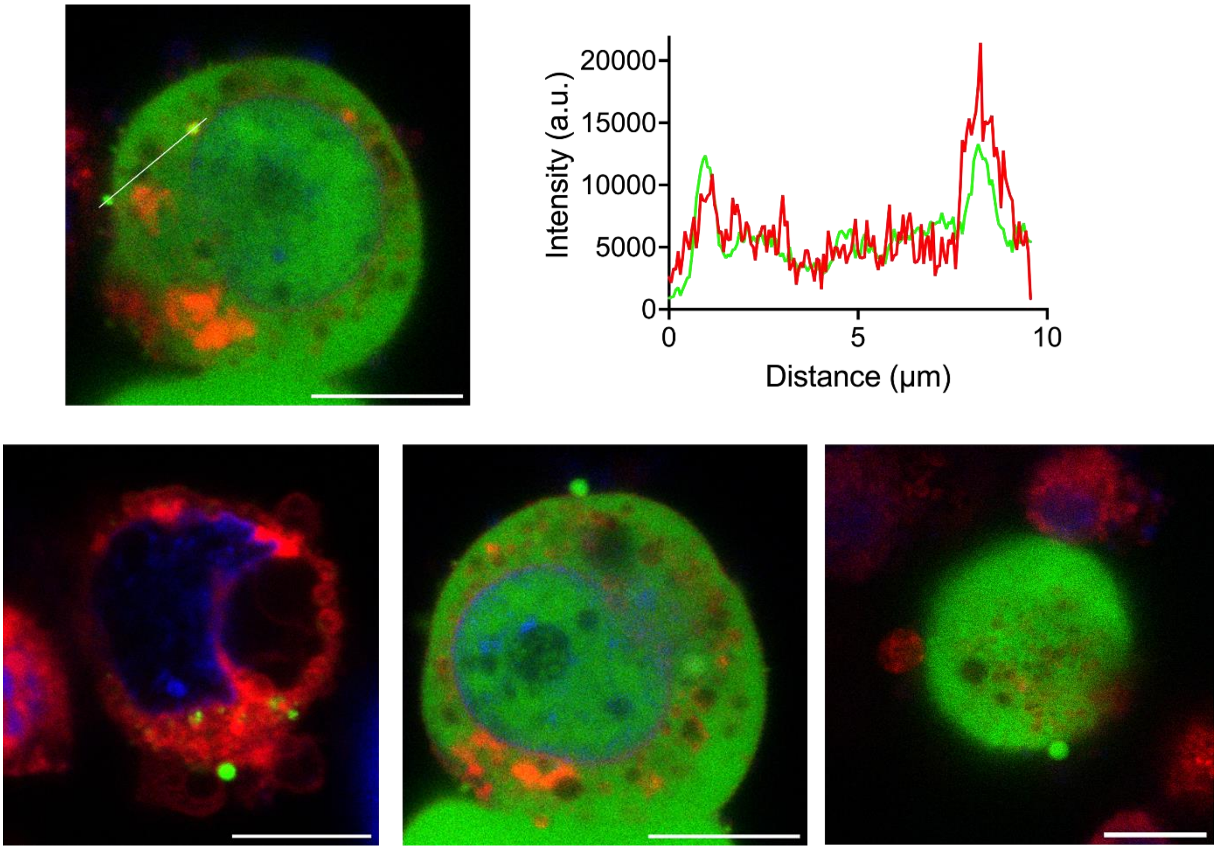

Supp figure 6: Top left: Confocal fluorescence 2D plane showing Xpa enriched vesicles in the cytoplasm and cell surface with the Xpa fluorescence (green) and ER tracker fluorescence (red) top right: line plot corresponding to the white line on the left image. Bottom: three examples of Xpa rich vesicles found on the surface of one low Xpa expressing cell (left) and two high Xpa expressing cells (right). Green channel: Xpa fluorescence, red channel ER tracker fluorescence, blue channel: hoechst fluorescence. All scale bars 10µm

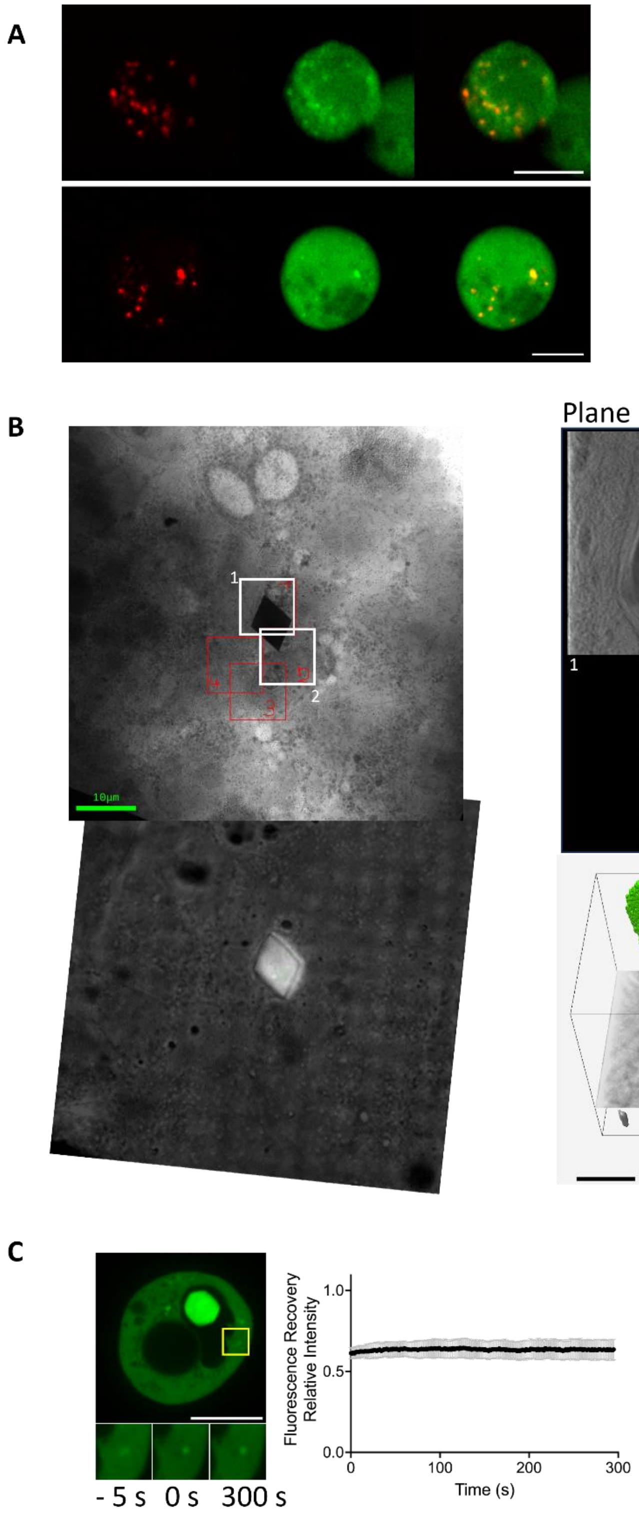

Supp figure 7: (A) snapshots of multiphoton (fluorescence, SHG and merged) from supp movies 5 and 6 showing colocalization of fluorescent and SHG puntae in the cytoplasm of two cells. Scale bars 10µm. (B) left: large cryoSXT images of a type I.C crystal in a cell (top) and corresponding oriented fluorescence imageof the crystal. On the cryoSXT image are represented the area chosen the 4 tomograms mosaic of high resolution imaging named 1 to 4. The two white squares delineate the tomograms represented on the right. Right top: two 2D planes of the reconstructed tomograms 1 and 2. The white arrows shows crystals engulphed in vesicles next to the large type I.C crystal. Right bottom: Volume segmentation of the area of tomogram 2 showing in green part of the large type I.C Xpa crystal and both Xpa crystals in vesicles. Grey structures : gold fiducials. Scale bars 1µm. (C) Fluorescence images with Xpa dots of a representative cell analyzed by FRAP. Scale bar: 10 µm. Quantification of the FRAP data from 6 different experiments.

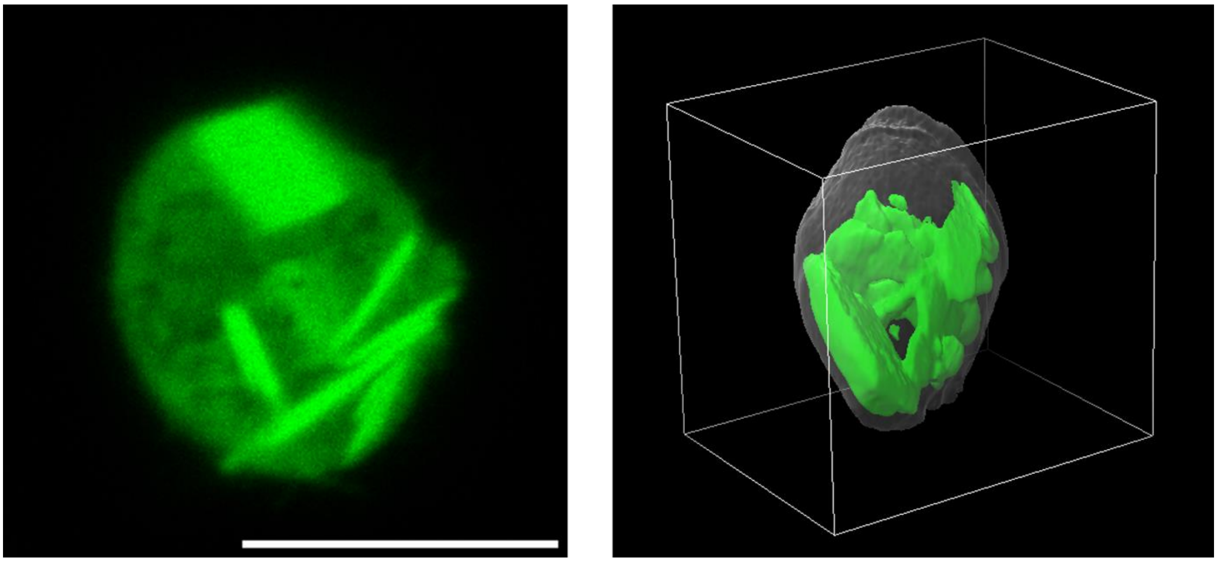

Supp figure 8: left: confocal fluorescence plane of a Xpa sorbitol induced plates only containing cell (scale bar 10µm). Right 3D segmentation of the confocal fluorescent volume acquired. green structures: Xpa plates, semi transparent white structure: outer cell surface.
